# *Arabidopsis* root-type ferredoxin:NADP(H) oxidoreductases are crucial for root growth and ferredoxin-dependent processes

**DOI:** 10.1101/2025.02.02.636142

**Authors:** Kota Monden, Daisuke Otomaru, Takamasa Suzuki, Tsuyoshi Nakagawa, Takushi Hachiya

**Affiliations:** Department of Molecular and Functional Genomics, Interdisciplinary Center for Science Research, Shimane University, 1060 Nishikawatsu-cho, Matsue, Shimane 690-8504, Japan; College of Bioscience and Biotechnology, Chubu University, 1200 Matsumoto-cho, Kasugai, Aichi 487-8501, Japan

**Keywords:** ferredoxin, ferredoxin:NADP(H) oxidoreductase, grafting, root growth, root-type ferredoxin:NADP(H) oxidoreductase, sulfur metabolism

## Abstract

Root-type ferredoxin:NADP(H) oxidoreductase (RFNR) is believed to reduce ferredoxin using NADPH in nonphotosynthetic tissues, facilitating ferredoxin-dependent biological processes. However, the physiological functions of RFNR remain unclear due to the difficulty in obtaining mutants lacking redundant RFNR isoproteins. The present study successfully generated *Arabidopsis* homozygous *rnfr1*;*2* double mutants by traditional crossing and selection. However, they displayed severely stunted roots, challenging subsequent growth and abundant seed recovery. Notably, grafted plants combining mutant scions with wild-type rootstocks exhibited normal growth and produced abundant mutant seeds. Growth analysis employing reciprocal grafts with the wild-type and mutant plants showed that primary root growth was inhibited only when the rootstock was derived from the mutants. Meanwhile, the absence of RFNR1 and 2 in the scion had no apparent impact on shoot and root growth. Root transcriptome analysis revealed that RFNR1 and 2 deficiency upregulated genes encoding ferredoxin-dependent enzymes and root-type ferredoxin, leading to genome-wide reprogramming associated with cell walls, lipids, photosynthesis, secondary metabolism, and biotic/abiotic stress responses. Thus, *Arabidopsis* RFNR1 and 2 are crucial for root growth and various ferredoxin-dependent biological processes.

## 1. Introduction

Ferredoxin:NADP(H) oxidoreductase (FNR) is central to plastidic redox metabolism in plants [1,2]. In chloroplasts, FNR oxidizes reduced ferredoxin to reduce NADP^+^, generating NADPH to drive the Calvin cycle, which is vital for autotrophic growth. In nonphotosynthetic plastids, FNR reduces ferredoxin (Fd) utilizing NADPH from the oxidative pentose phosphate pathway (oxPPP). Reduced Fd can supply reducing power to the enzymes involved in nitrite and sulfite reduction, glutamate biosynthesis, monodehydroascorbate reduction, and fatty acid desaturation [2,3,4]. These opposing reactions catalyzed by FNR are regulated by two distinct isoforms: leaf-type FNR (LFNR) in chloroplasts and root-type FNR (RFNR) in nonphotosynthetic plastids, each exhibiting varying kinetic properties, redox potentials, spatio-temporal expression patterns, and three-dimensional structures [1,5,6,7,8,9]. Moreover, multiple Fd isoforms allow selective interactions with FNR isoforms, which enables elaborate electron channeling depending on the cell type and environment [1,2,8,10,11].

*Arabidopsis* possess two *LFNR*s, *AtLFNR1* and *AtLFNR2;* and *RFNR*s, *AtRFNR1* and *AtRFNR2* [1]. Deficiency of either *AtLFNR1* or *AtLFNR2* reduces rosette size, pales green leaves, and decreases CO_2_ fixation, while PSII activity remains mostly unchanged [12,13]. The double mutant of *AtLFNR1* and *AtLFNR2* possesses only a few malformed chloroplasts with unstacked thylakoid membranes and cannot grow under autotrophic conditions without sucrose supplementation [14]. Thus, AtLFNR1 and 2 function redundantly to sustain photosynthesis *in planta*. Meanwhile, expression profiling suggests that AtRFNR1 and 2 function mainly in the roots, with AtRFNR2 being the major isoform [7,9]. The transcriptional induction of *AtRFNR1* by *AtRFNR2*-knockout and the strong coexpression of *AtRFNR1* and *2* implies their redundant roles [7]. When recombinant *Arabidopsis* Fd was reduced by recombinant maize FNR employing NADPH *in vitro*, maize LFNR exhibited comparable kinetic parameters for *Arabidopsis* leaf-type (AtFd1 and AtFd2) and root-type Fd (AtFd3); maize RFNR had a higher affinity for AtFd3 than AtFd1 or AtFd2 [6]. Moreover, an *in vitro* reduction system comprising NADPH, maize RFNR, and AtFd demonstrated that AtFd3 showed greater activity and affinity for sulfite reductase (SiR) compared with AtFd1 and AtFd2 [6]. These findings suggest that AtRFNRs contribute to Fd-dependent biosynthetic processes by reducing AtFd3. However, previous studies have failed to identify any visible phenotype in plants lacking either AtRFNR1 or 2 in commonly used media or soils [7,9], except nitrite as the sole N source [7]. Unfortunately, to date, homozygous mutants lacking both AtRFNR1 and 2 have not been obtained [9]. The lethality of the double mutant is likely due to defects in gamete rather than embryo development, suggested by the progeny segregation ratio and the absence of aborted seeds in the parent silique [9]. Consequently, the physiological functions of RFNR remain unclear.

In the present study, we generated *Arabidopsis* homozygous *rnfr1*;*2* double mutants from parental lines used previously [9]. They displayed severely stunted root growth, challenging subsequent growth and abundant seed recovery. Notably, we discovered that grafted plants combining mutant scions with wild-type rootstocks exhibited normal growth and produced abundant mutant seeds. Our growth and transcriptome analyses indicated that AtRFNR1 and 2 are crucial for root growth and various Fd-dependent biological processes *in planta*.

## 2. Materials and Methods

### 2.1. Plant materials

In this study, we used *A. thaliana* accession Col-0 (Col) as the control line as well as the *rfnr1* (SALK_085009), *rfnr2-1* (SAIL_527_G10), and *rfnr2-2* (SALK_133654) mutants [7,9]. The *rfnr1*;*rfnr2-1* and *rfnr1*;*rfnr2-2* double mutants were generated via crossing.

### 2.2. Plant growth analysis

For *in vitro* culture, the surface-sterilized seeds were placed on a solid medium containing 50 mL half-MS salts without vitamins (Code M0221; Duchefa Biochemie, Haarlem, Netherlands) supplemented with 0.05% (w/v) MES-KOH (pH 5.7), 1% (w/v) sucrose, and 0.5% (w/v) gellan gum (Fujifilm Wako, Osaka, Japan). The plants were grown vertically under a PPFD of 25 μmol m^−2^ s^−1^ (constant light) at 22°C. The plates were scanned at 600 dpi with a GT-X980 instrument (EPSON, Tokyo, Japan), and the root length was measured using the ImageJ software version 1.52p (https://imagej.net/ij/). For soil cultivation, the plants were grown under a PPFD of 25–35 μmol m^−2^ s^−1^; 16/8-h light/dark cycle; at 19°C–20°C on Jiffy-7 peat pellets (Jiffy Products International AS, Kristiansand, Norway) or on Supermix A peat soil (Sakata Seed, Kyoto, Japan).

### 2.3. Micrografting

Micrografting followed a previously described method [15] with slight modifications. The seedlings were grown vertically on a solid medium containing 50 mL of half-MS salts including vitamins (Code M0222; Duchefa Biochemie) supplemented with 0.05% (w/v) MES-KOH (pH 5.7), 1% (w/v) sucrose, and 0.5% (w/v) gellan gum (Fujifilm Wako, VA, USA) under a PPFD of 25 μmol m^−2^ s^−1^; 16/8-h light/dark cycle at 22°C. The 4-day-old seedlings were cut perpendicularly in the hypocotyl with an injection needle tip (Code NN-2613S; TERUMO, Tokyo, Japan). The obtained scion was connected with the partner rootstock in a silicon microtube (Code 1-8194-04; AS ONE, Osaka, Japan). The grafted plants were incubated for 5 days under a PPFD of 50 μmol m^−2^ s^−1^ (constant light) at 27°C. Successfully grafted plants without adventitious root formation were selected and used in the subsequent experiments.

### 2.4. Growth analysis of the grafted plants

For *in vitro* culture, the grafted plants were transferred to a solid medium containing 50 mL of half-MS salts without vitamins (Code M0221; Duchefa Biochemie) supplemented with 0.05% (w/v) MES-KOH (pH 5.7), 1% (w/v) sucrose, and 0.5% (w/v) gellan gum (Fujifilm Wako), and grown vertically for 3 days under a PPFD of 25 μmol m^−2^ s^−1^ (constant light) at 22°C. The plates were scanned at 600 dpi (GT-X980; EPSON), and the root length was measured with ImageJ version 1.52p. For soil cultivation, the grafted plants were transferred onto Jiffy-7 peat pellets and grown for 23 days under a PPFD of 100 μmol m^−2^ s^−1^;16/8-h light/dark cycle; at 22°C–23°C.

### 2.5. Total protein determination

The seedlings were grown vertically for 4 days on a solid medium containing 50 mL of half-MS salts without vitamins (Code M0221; Duchefa Biochemie) supplemented with 0.05% (w/v) MES-KOH (pH 5.7), 1% (w/v) sucrose, and 0.5% (w/v) gellan gum (Fujifilm Wako) under a PPFD of 25 μmol m^−2^ s^−1^ (constant light) at 22°C. The excised roots from 100 seedlings were flash-frozen in liquid N_2_ as one biological replicate. The frozen roots were homogenized with a multi-bead shocker (Yasui Kikai) employing zirconia beads. Total proteins were extracted with sample buffer (2% [w/v] SDS, 62.5 mM Tris-HCl [pH 6.8], 10% [v/v] glycerol, and 0.0125% [w/v] bromophenol blue) supplemented with Halt^TM^ protease inhibitor cocktail (Thermo Fisher Scientific, Tokyo, Japan), and incubated at 95°C for 5 min. The extracts were centrifuged at 20,400× g at 8°C for 10 min. The protein concentration of the supernatant was determined with a Qubit™ Protein Broad Range Assay Kit (Thermo Fisher).

### 2.6. Immunodetection of RFNR proteins

SDS-PAGE and immunodetection were per a previously described method [16] with slight modifications. The protein extract was mixed with dithiothreitol and incubated at 95°C for 3 min. A 7 μL-mixture containing 15.3 μg of protein per lane was subjected to SDS-PAGE using a 12% Mini-PROTEAN TGX Gel (Bio-Rad, CA, USA) and transferred to a Trans-Blot Turbo Mini PVDF membrane utilizing the Trans-Blot Turbo Transfer System (Bio-Rad). The membranes were incubated for 1 h in blocking buffer (50 mM Tris-HCl and 150 mM NaCl, pH 7.6, 0.1% [v/v] Tween-20) containing 5% (w/v) ECL Prime Blocking Agent (Cytiva, Tokyo, Japan). The membranes were incubated for 1 h with a polyclonal antibody at 1:50,000 dilution raised against maize RFNR [10]. After rinsing, the antigen–antibody complex was detected with a horseradish peroxidase-conjugated antirabbit IgG donkey antibody (Cytiva) at 1:50,000 dilution and visualized ECL Prime chemiluminescent detection (Cytiva). Proteins blotted on the membranes were visualized using the CBB Protein Safe Stain (TaKaRa). Chemiluminescence and CBB staining were captured with an ImageQuant LAS 500 (Cytiva).

### 2.7. RNA extraction

The excised roots from 200 seedlings were grown under the same conditions as section 2.5. They were flash-frozen in liquid N_2_ as one biological replicate. The frozen roots were homogenized using zirconia beads in a multi-bead shocker (Yasui Kikai). Total RNA was extracted using an RNeasy Plant Mini Kit (Qiagen, Tokyo, Japan) with on-column DNase digestion (Qiagen).

### 2.8. RNA-seq

RNA quality was evaluated with a Qubit RNA IQ Assay Kit (Thermo Fisher Scientific). RNA samples with IQs of 9.8–10.0 were utilized for library preparation. cDNA libraries were constructed employing a NEBNext Ultra II RNA Library Prep Kit with Sample Purification Beads, an NEBNext Poly(A) mRNA Magnetic Isolation Module, and NEBNext Multiplex Oligos (New England Biolabs, Tokyo, Japan) for Illumina. cDNA libraries were sequenced on a NextSeq 500 platform (Illumina, Tokyo, Japan), and the resulting bcl files were converted to fasta format using bcl2fastq (Illumina). The raw reads were mapped to the TAIR10 by Bowtie [17]. The read counts thus obtained were analyzed utilizing iDEP version 096 [18], Metascape [19], and MapMan [20].

### 2.9. RT-qPCR

Reverse transcription (RT) employed a ReverTraAce qPCR RT Master Mix with gDNA Remover (Toyobo, Osaka Japan). The synthesized cDNA was diluted 10-fold with water and used for qPCR on a QuantStudio 1 instrument (Thermo Fisher Scientific) with KOD SYBR qPCR Mix (Toyobo, Osaka, Japan). Relative transcript levels were calculated by applying the comparative cycle threshold method with *TIP41* (AT4G34270) as an internal standard [21]. The primer sequences are shown in Supplementary Table 1.

### 2.10. Statistical analysis

The Tukey-Kramer multiple comparison and χ^2^ tests were performed using R software v.2.15.3 (https://cran.r-project.org/bin/windows/base/old/). For the χ^2^ test, the expected segregation ratio for the genotypes *RFNR1 RFNR2* (+/+) (+/+), (+/+) (+/−), (+/+) (−/−), (+/−) (+/+), (+/−) (+/−), (+/−) (−/−), (−/−) (+/+), (−/−) (+/−), and (−/−) (−/−) was set at 1:2:1:2:4:2:1:2:1, respectively.

## 3. Results and Discussion

### 3.1. Recovery of the RFNR1;2-double mutant seeds using the micrografting technique

Our examination of the *rfnr1* × *rfnr2-1* and *rfnr1* × *rfnr2-2* F_2_ progeny identified several plants with severely stunted root growth (Fig. 1A). Genomic PCR confirmed them as homozygous T-DNA insertion mutants for *RFNR1* and *2*, i.e., *rfnr1*;*rfnr2-1* (DM1) and *rfnr1*;*rfnr2-2* (DM2) (Figs. 1B and C). However, severe growth impairment in these double mutants rendered seed recovery challenging (Fig. 1A and Supplementary Fig. 1). Because RFNR1 and 2 are expressed predominantly in the roots [7,9], the severe growth phenotype of the mutants may be caused by RFNR1 and 2 deficiency in the roots. Then, we grafted the mutant scions onto the Col rootstocks employing our micrografting technique. Strikingly, the grafted plants obtained DM1/Col and DM2/Col (scion/rootstock) exhibited normal vegetative growth comparable to the control Col/Col (Fig. 1D), allowing the recovery of abundant DM1 and DM2 seeds for further experiments (Fig. 1E). RT-PCR and immunoblots of DM2 roots did not detect any signals corresponding to *RFNR1* and *2* and RFNR1 and 2, respectively (Figs. 1F and G). These findings support our previous report that *rfnr1* and *rfnr2-2* are knockout alleles [7]. Meanwhile, the *RFNR2* signal was lower in DM1 than in Col when employing primers spanning exons 1– 4 (Figs. 1B and F), confirming that *rfnr2-1* is a knockdown allele as was reported [7]. However, primers spanning the 5′-UTR to the fourth exon failed to detect *RFNR2* in DM1 (Figs. 1B and F). To resolve this contradiction, we performed RNA-seq and mapped the obtained reads to each splice variant of *RFNR2* (Supplementary Fig. 2). The reads aligned to the upstream region of the 5′-UTR of the representative model *AT1G30510.2* and its counterpart *AT1G30510.1* in Col, but not in DM1. Given that RFNR1 and 2 signals were not detected in DM1 (Fig. 1G), it is plausible that *RFNR2*, lacking the 5′-UTR, may not be translated in DM1. Meanwhile, the read distribution for *AT1G30510.3*, which encodes a smaller isoprotein, was similar between Col and DM1, suggesting the absence of AT1G30510.3 proteins in both lines. In summary, our micrografting technique produced abundant seeds of mutants deficient in both AtRFNR1 and 2.

**Fig. 1.**
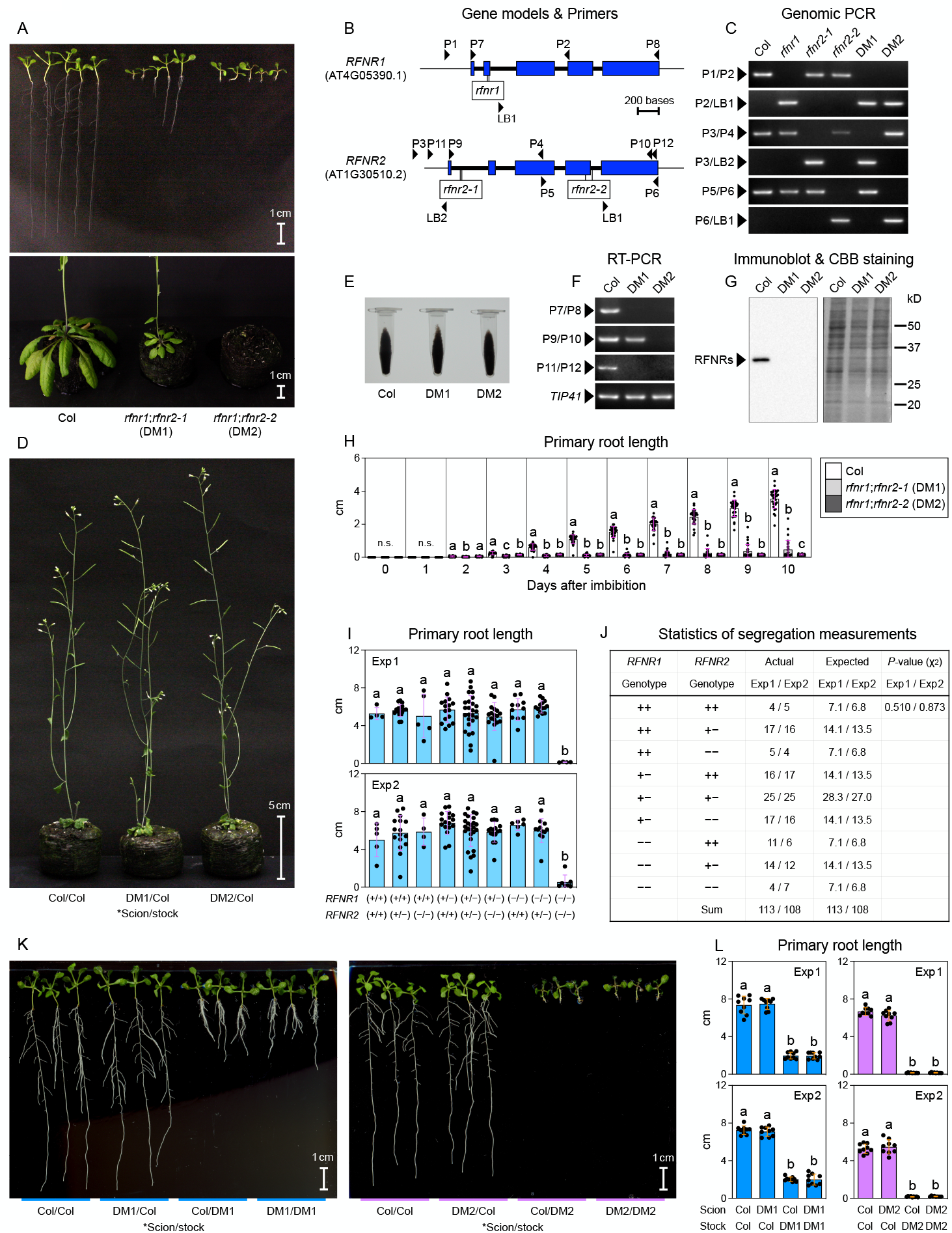
Root-expressed AtRFNRs are vital for primary root growth. (A) The appearance of 10-day-old seedlings grown on half-MS medium (upper) and 56-day-old plants grown on Jiffy-7 peat pellets (lower). (B) Schematic representative gene models for *RFNR1* (AT4G05390.1) and *RFNR2* (AT1G30510.2) along with the positions of the primers used for genomic-(C) and RT-PCR (F). The scale bar represents 200 bases. (C) Genomic PCR and agarose gel electrophoresis results. (D, E) The appearance of 32-day-old grafted plants grown on Jiffy-7 peat pellets (D) and the seeds harvested from the grafted plants carrying scion-derived genotypes (E). (F) RT-PCR and agarose gel electrophoresis results. (G) Results of immunoblotting employing the polyclonal antibody raised against maize RFNR and Coomassie Brilliant Blue staining. The arrowhead indicates the signals corresponding to RFNR1 and 2. (H) Time-course of the primary root length after imbibition and grown on half-MS medium. (I, J) Primary root length of 12-day-old F_2_ seedlings derived from *rfnr1* and *rfnr2-1* grown on half-MS medium (I), along with the results of the χ^2^ test applied for evaluating the Mendelian segregation ratio in these seedlings (J). (K, L) The appearance of 12-day-old grafted seedlings grown on half-MS medium (K) and their primary root length (L). (H, I, L) Various lowercase letters indicate statistically significant differences as determined by the Tukey-Kramer tests at *P* < 0.05. Data: mean ± SD; n = 30 (H), n = 4–25 (I), n = 9 (L). n.s. denotes not significant. (I, J, L) The results from two independent experiments, i.e., Exp1 and Exp2, are shown.

### 3.2. Root-expressed AtRFNRs are vital for primary root growth

We mapped the time course of primary root lengths to examine the effect of RFNR1 and 2 on root growth (Fig. 1H). From the third day after imbibition, the lengths of DM1 and DM2 remained markedly shorter than that of Col. Interestingly, in some DM1 individuals, primary root elongation continued after the third day, whereas, it almost ceased in DM2. The primary roots of DM1 were significantly longer than those of DM2, ten days after imbibition. This result suggests that DM1 is a slightly weaker allele than DM2, although the reasons remain unclear.

Next, to reveal the effects of *RFNR1* and *RFNR2* interactions on primary root growth, we measured the primary root length of 12-day-old F_2_ seedlings derived from *rfnr1* and *rfnr2-1* (Fig. 1I). Primary root elongation was impaired only in the seedlings lacking both *RFNR1* and *2*, indicating the redundant roles of RFNR1 and 2 in primary root growth. Notably, the *P* values from the χ^2^ test utilizing the Mendelian segregation ratio of F_2_ seedlings were much higher than 0.05 (Fig. 1J). This finding suggested that viable F_2_ progeny were obtained per Mendelian laws, thereby rejecting the possibility of gametophyte lethality resulting from a deficiency in both *RFNR1* and *2* as suggested previously [9], at least under our growth conditions.

Furthermore, to clarify whether shoot- or root-expressed RFNR1 and 2 affect primary root growth, we determined the primary root length employing reciprocally grafted seedlings (Figs. 1K and L). Irrespective of the scion’s genotype, the primary root growth was severely inhibited only when the rootstock was derived from DM1 or DM2, more in DM2 than in DM1, corresponding to the observation reported in Figure 1H. Meanwhile, the absence of RFNR1 and 2 in the scion had no apparent impact on shoot and root growth (Figs. 1D and K). In summary, root-expressed RFNR1 and 2 are vital for primary root growth.

### 3.3. RFNR is crucial for ferredoxin-dependent processes in the root

To overview the roles of RFNRs in the root, we performed RNA-seq analysis using roots from 4-day-old seedlings. The raw read counts, reads per million mapped reads, and normalized transcript levels are presented in Supplementary Tables 2 and 3. Hierarchical clustering employing iDEP [18] classified the transcripts into two groups depending on the presence or absence of RFNR1 and 2 (Fig. 2A). Therefore, the differentially expressed genes (DEGs) between Col and DM1/DM2 were explored (Fig. 2B and Supplementary Table 4). An analysis of the DEGs with a minimum fold-change of 2 revealed that 2,436 genes were significantly upregulated in both DM1 and DM2 relative to Col-0, whereas 2,120 were downregulated. Even at a minimum fold-change of 10, 632 and 234 genes were up and downregulated by RFNR1 and 2 deficiency, respectively, indicating an impactful role of RFNR1 and 2 in root activity. Notably, RFNR1 and 2 deficiency induced genes encoding the plastidic Fd-dependent enzymes involved in S assimilation (*SIR*), N assimilation (*NIR* and *GLU1*), monodehydroascorbate reduction (*MDAR5*), and fatty acid desaturation (*FAD5/6/7/8*) (Figs. 2C, D and Supplementary Fig. 3) [2,3,4]. Additionally, the expression of genes encoding root-type Fd (*FD3*) and plastidic dehydrogenases (*G6PD2, G6PD3*, and *PGD3*) that produce NADPH for Fd reduction in the oxPPP exhibited similar responses (Figs. 2E and F). These inductions are considered a compensatory response to depletion of reduced Fd. Moreover, enrichment analysis of DEGs with a minimum fold-change of 10 (Supplementary Fig. 4) and profiling of metabolism-related genes (Supplementary Fig. 5) revealed that genome-wide reprogramming occurred due to RFNR1 and 2 deficiency, especially in genes associated with the cell wall, lipids, secondary metabolism, photosynthesis, and biotic/abiotic stress responses. This observation suggests that RFNR1 and 2 could sustain a wide range of biological processes by maintaining redox homeostasis in the roots.

**Fig. 2.**
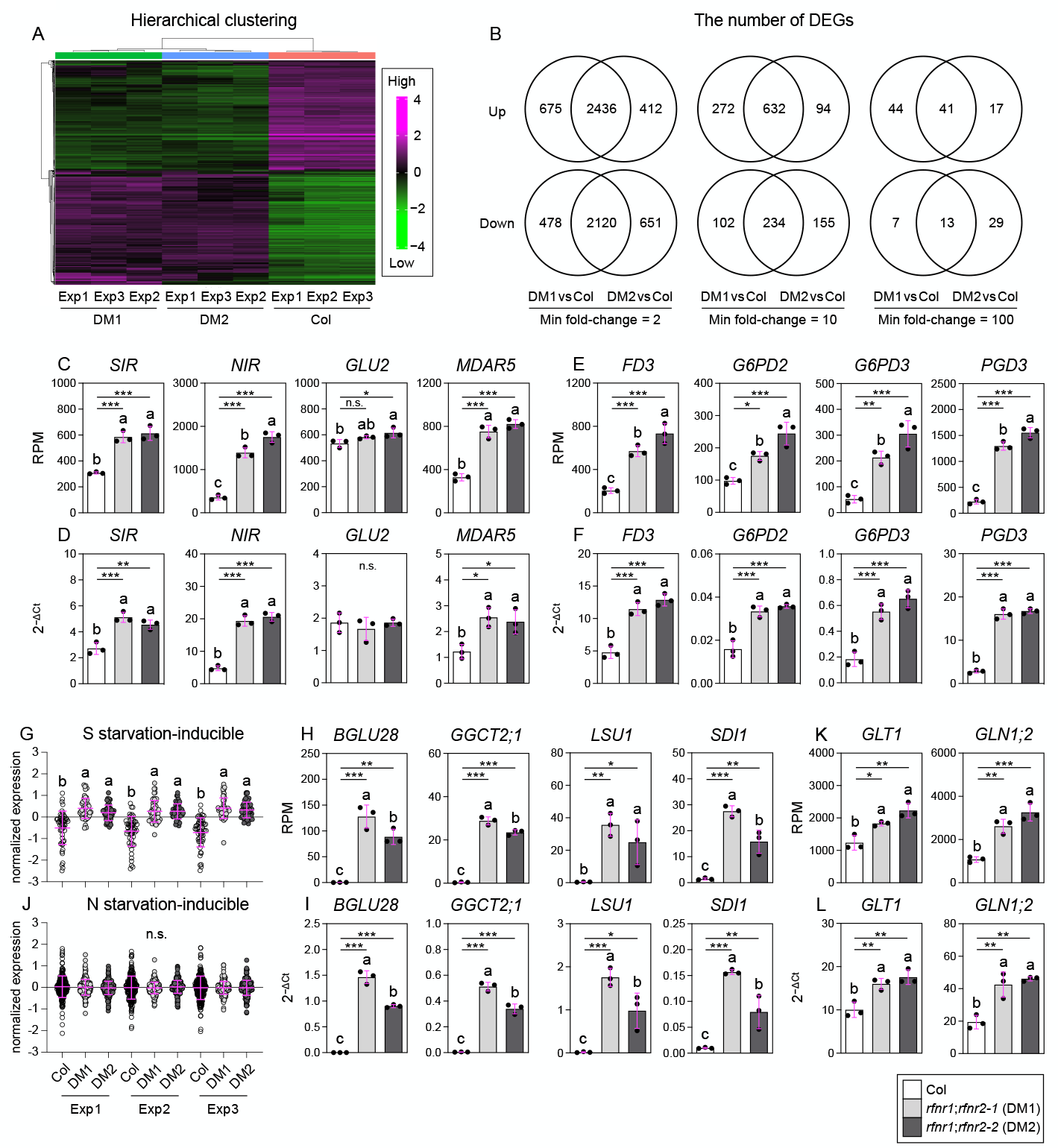
AtRFNRs are crucial for ferredoxin-dependent processes in the roots. (A) Heat map generated from the hierarchical clustering of normalized transcript levels. Magenta and green represent higher and lower expression levels, respectively. (B) Venn diagram showing the number of DEGs in DM1 and/or DM2 versus Col, with minimum fold-change thresholds of 2, 10, and 100. (C, D) The expression of root genes encoding plastidic Fd-dependent enzymes, as determined by RNA-seq (C) and RT-qPCR (D). (E, F) The expression of genes encoding root-type Fd and plastidic dehydrogenases in the oxPPP, as determined by RNA-seq (E) and RT-qPCR (F). (G, J) Root expression of S-(G) and N-starvation-inducible genes (J), as determined by RNA-seq. Data: mean ± SD. The gene lists were obtained from previous studies [22,23]. (H, I) Expression of the root S-starvation-inducible genes, as determined by RNA-seq (H) and RT-qPCR (I). (K, L) The expression of genes encoding enzymes vital for primary ammonium assimilation in the root, as determined by RNA-seq (K) and RT-qPCR (L). (A, G, J) Exp 1, 2, and 3 denote the first, second, and third experiments, respectively. (C–L) Various lowercase letters indicate statistically significant differences ascertained by Tukey-Kramer tests at *P* < 0.05 (\**P* < 0.05; \*\**P* < 0.01; \*\*\**P* < 0.001). n.s. denotes not significant. (C–F, H, I, K, and L) Data = mean ± SD (n = 3).

Because sulfite reductase, nitrite reductase, and Fd-dependent glutamate synthase receive electrons from reduced Fd [2], RFNR1 and 2 deficiency may impair S and N assimilation, causing starvation. As expected, S starvation-inducible genes were markedly upregulated by this deficiency (Fig. 2G and Supplementary Table 5). Several S-starvation-associated markers (*BGLU28, GGCT2;1, LSU1*, and *SDI1*) [22] were upregulated extremely highly (Figs. 2H and I), with S transporter-encoding genes exhibiting the most consistent upregulation among the various transporter- and channel-encoding genes (Supplementary Fig. 6). Thus, our observations suggest that RFNR1 and 2 are vital for S metabolism in the roots. Meanwhile, N-starvation-inducible genes [23] were not significantly upregulated by this deficiency (Fig. 2J and Supplementary Table 5). Notably, ammonium assimilation bypasses the Fd-dependent nitrite reaction catalyzed by NIR and can be primarily driven by glutamine synthase GLN1;2 and the NADH-dependent glutamate synthase GLT1 in *Arabidopsis* roots [24,25]. Since our growth media included nitrate and ammonium as N sources, the induction of *GLN1;2* and *GLT1* by RFNR1 and 2 deficiency (Figs. 2K and L) may enhance ammonium assimilation and compensate for the lack of nitrate utilization, thereby avoiding N starvation.

### 3.4. Conclusion

We obtained the abundant seeds of *Arabidopsis* mutants lacking the RFNR isoproteins through our micrografting technique. Growth and transcriptome analysis of the mutants revealed that RFNR1 and 2 are important for root growth and various Fd-dependent processes. Future metabolomic and reductant profiling of the mutants may elucidate the precise physiological roles of RFNR1 and 2 and potentially discover novel Fd-dependent processes.

## Acknowledgments

The authors acknowledge Prof. Toshiharu Hase (Osaka University, Japan) for generously providing the RFNR antibodies. They also thank Mr. Takuto Kamikubo (Shimane University, Japan) for his advice concerning the experimental techniques.

## Author contributions: CRediT

Kota Monden: Writing-review and editing, Investigation, Fund acquisition. Daisuke Otomaru: Writing, review and editing, Investigation. Takamasa Suzuki: Writing, review and editing, Investigation. Tsuyoshi Nakagawa: Writing, review and editing, Supervision. Takushi Hachiya: Writing-original draft, Conceptualization, Validation, Formal analysis, Visualization, Supervision, and Fund acquisition.

## Data availability statement

The RNA-seq raw data are available in ArrayExpress (https://www.ebi.ac.uk/biostudies/arrayexpress) under accession number E-MTAB-14802.

