## Supplementary Fig. 1-6 for "*Arabidopsis* root-type ferredoxin:NADP(H) oxidoreductases are crucial for root growth and ferredoxin-dependent processes"

### Supplementary Figure 1

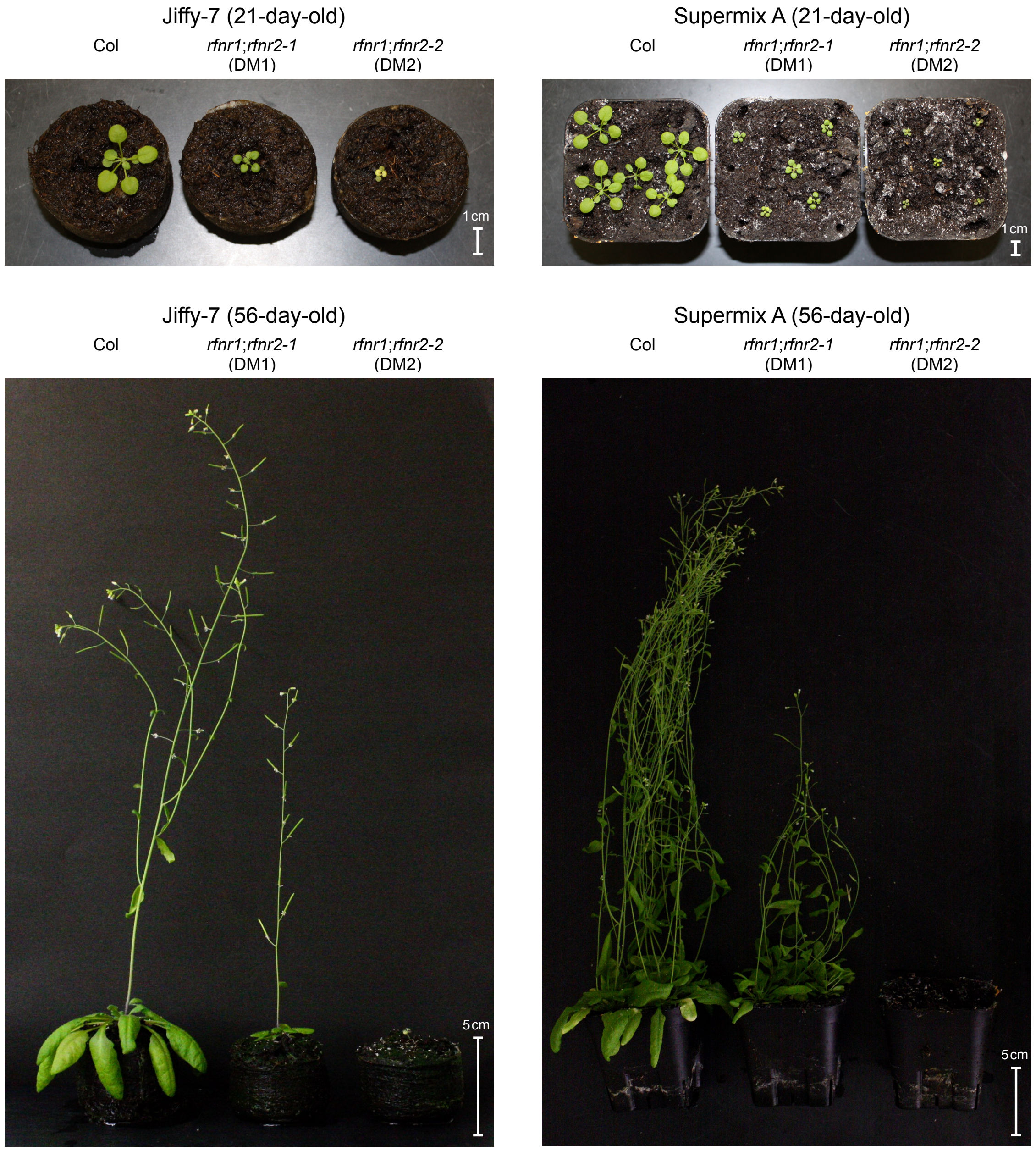


**Fig. S1. Representative photos of plant growth on Jiffy-7 peat pellets and on Supermix A peat soil.**

### Supplementary Figure 2

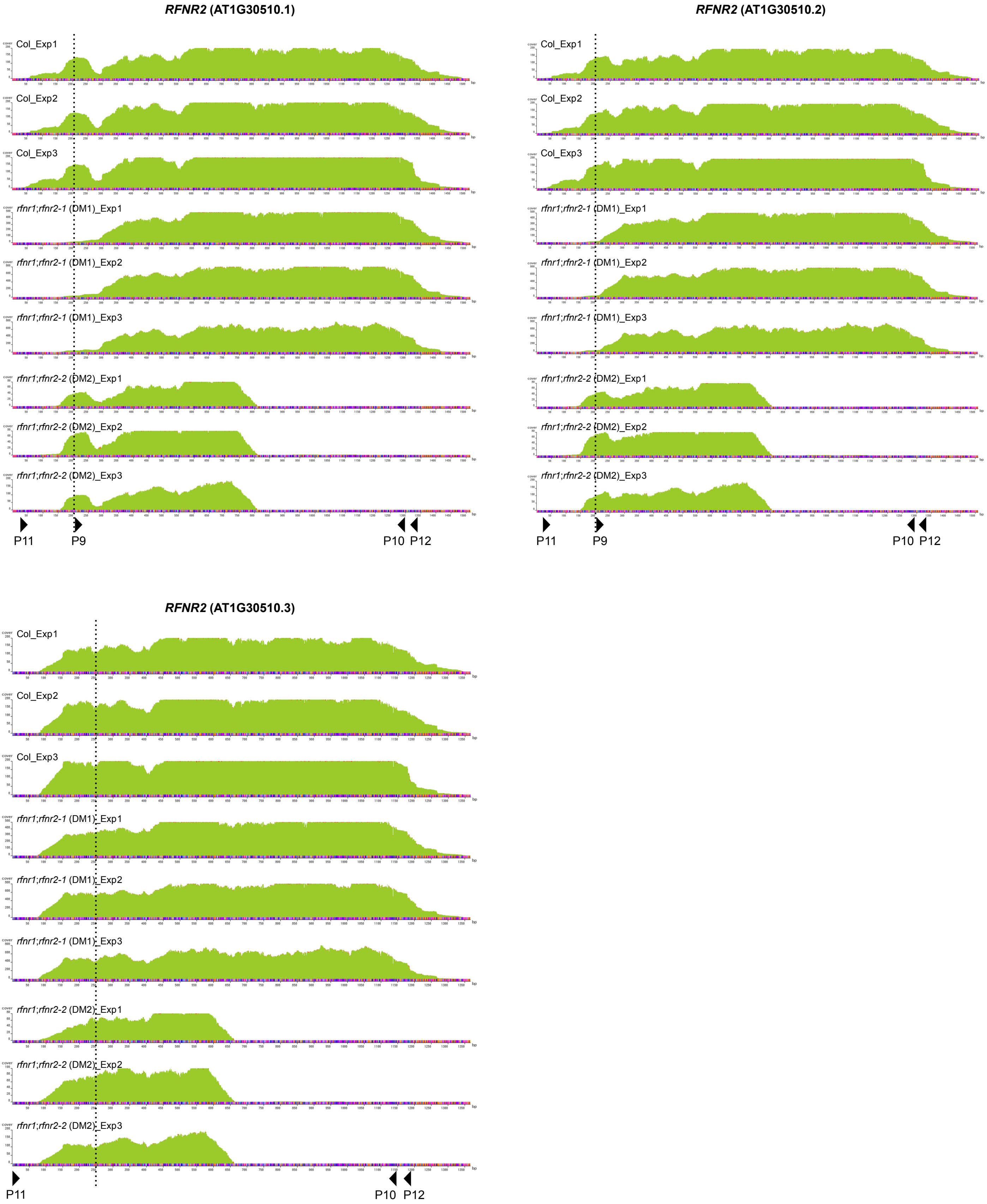


**Fig. S2. Mapping results of reads corresponding to each sprice variant of *RFNR2*.**
The dotted lines and arrow heads indicate the positions of the start codon and the specific primers, respectively. Note that AT1G30510.2 is the representative gene model in the TAIR database (https://www.arabidopsis.org/locus?key=29610).

### Supplementary Figure 3

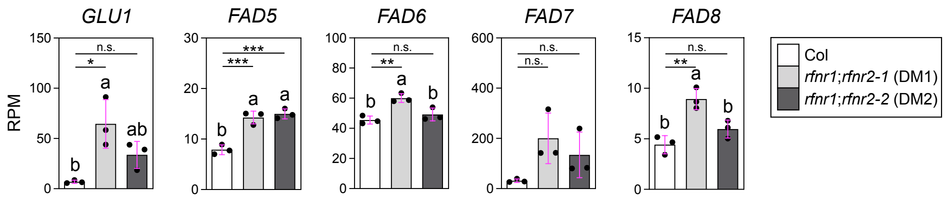


**Fig. S3. RPM values of plastidic glutamate synthase and fatty acid desaturase genes.**

### Supplementary Figure 4

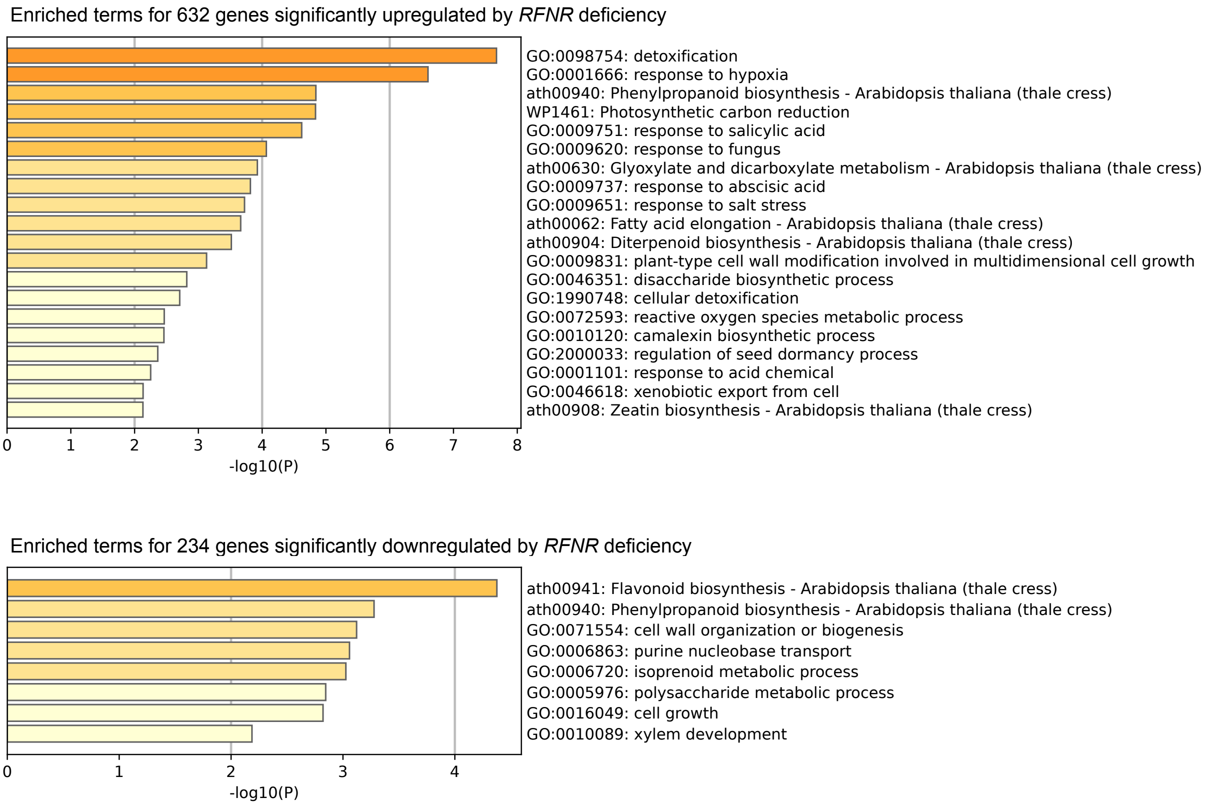


**Fig. S4. Enriched terms ourput from Metascape using DEGs with a minimum fold change of 10 as queries.** A list of DEGs is shown in Table S4.

### Supplementary Figure 5

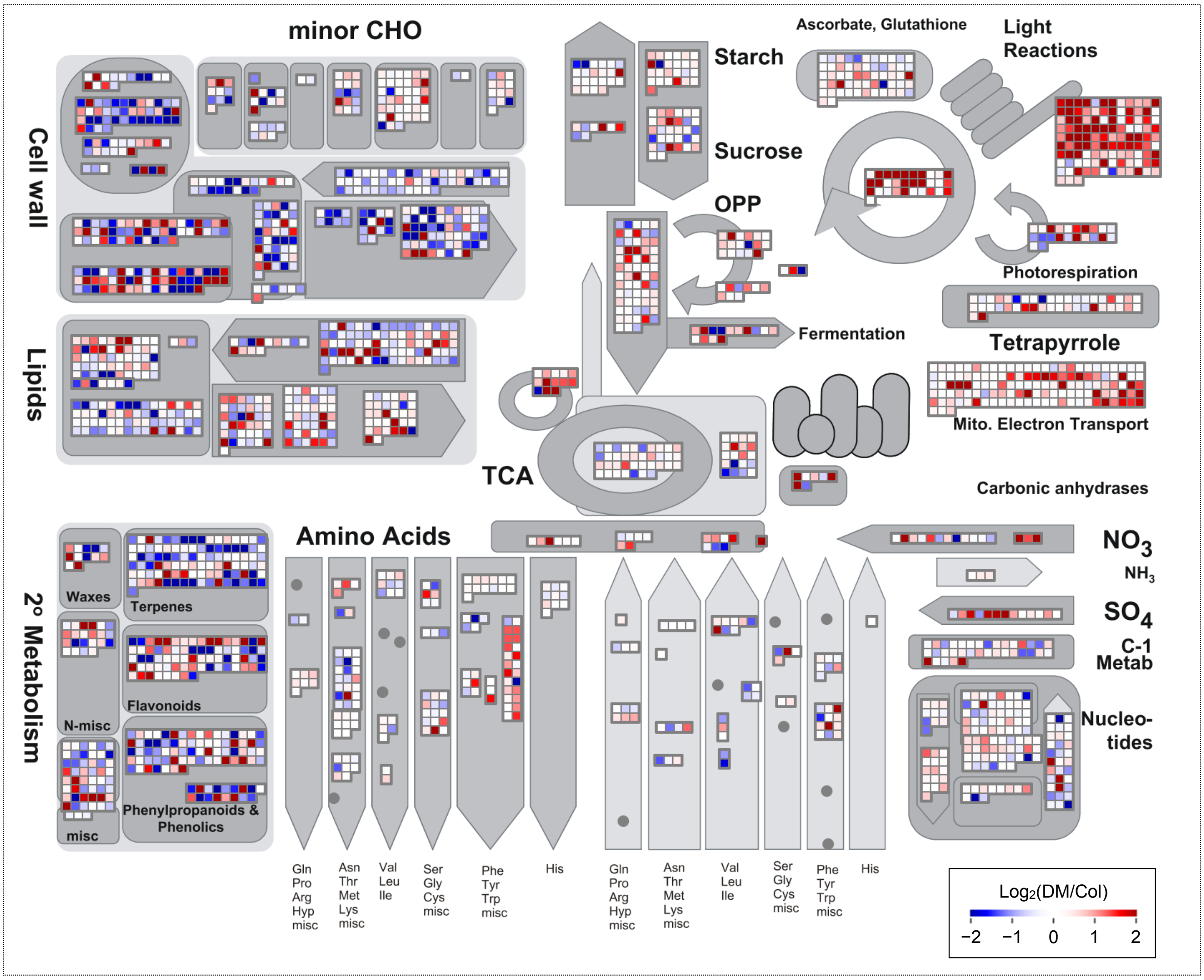


**Fig. S5. Outputs from MapMan ver 3.6.0RC1 on metabolism-related genes.**
For MapMan analysis, mean RPM values of DM1 and DM2 were utilized, and the values from three independent experiments were also averaged. The “Metabolism_overview” pathway (https://mapman.gabipd.org/mapmanstore) was employed for the analysis.

### Supplementary Figure 6

A

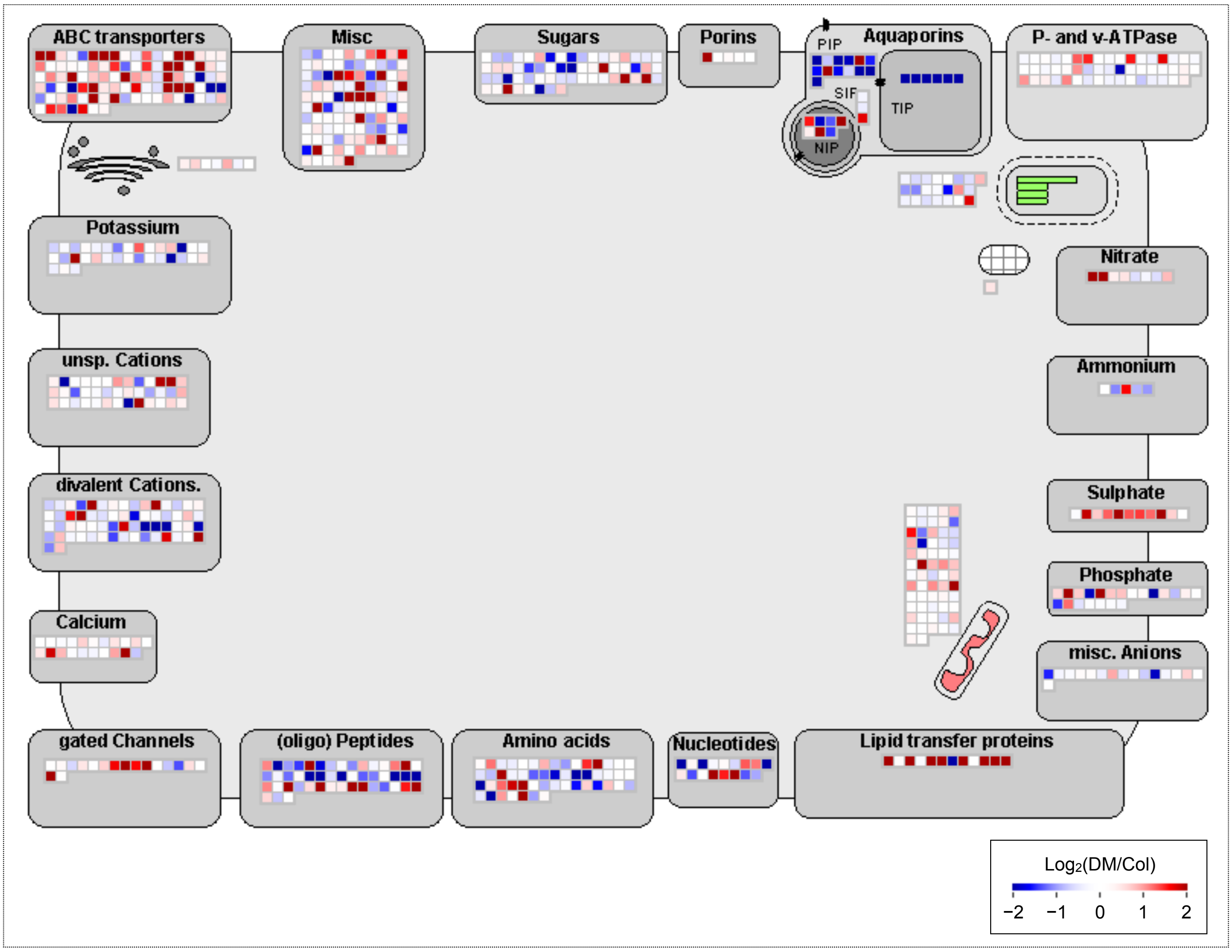


B

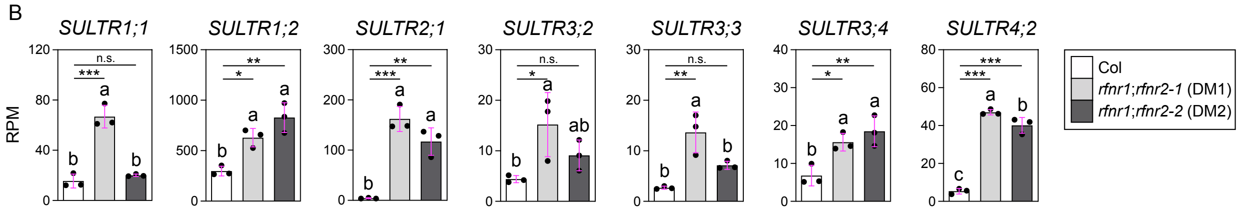


**Fig. S6. Outputs from MapMan ver 3.6.0RC1 on transporter and channel genes.**
(A) For MapMan analysis, mean RPM values of DM1 and DM2 were utilized, and the values from three independent experiments were also averaged. The "Transport_overview" pathway (https://mapman.gabipd.org/mapmanstore) was employed for the analysis. (B) RPM values of selected sulfate transporter genes.
